# MUSTANG: MUlti-sample Spatial Transcriptomics data ANalysis with cross-sample transcriptional similarity Guidance

**DOI:** 10.1101/2023.09.08.556895

**Authors:** Seyednami Niyakan, Jianting Sheng, Yuliang Cao, Xiang Zhang, Zhan Xu, Ling Wu, Stephen T.C. Wong, Xiaoning Qian

## Abstract

Spatially resolved transcriptomics has revolutionized genome-scale transcriptomic profiling by providing high-resolution characterization of transcriptional patterns. We here present our spatial transcriptomics analysis framework, **MUSTANG** (**MU**lti-sample **S**patial **T**ranscriptomics data **AN**alysis with cross-sample transcriptional similarity **G**uidance), which is capable of performing multi-sample spatial transcriptomics spot cellular deconvolution by allowing both cross-sample expression based similarity information sharing as well as spatial correlation in gene expression patterns within samples. Experiments on two real-world spatial transcriptomics datasets demonstrate the effectiveness of **MUSTANG** in revealing biological insights inherent in cellular characterization of tissue samples under the study. MUSTANG is publicly available at at https://github.com/namini94/**MUSTANG**

## 1. Introduction

Recent advances in single-cell **RNA** sequencing (sc**RNA**-seq) have enhanced our knowledge of different cellular development processes and can help better characterize heterogeneity of cell types in many complex tissues (Hwang *et al*., 2018; Niyakan *et al*., 2021). **H**owever, in original sc**RNA**-seq approaches spatial information is not retained when preparing samples with tissue dissociation and cell isolation Zhao *et al*. (2021). Thus, sc**RNA**-seq technologies lack the spatial resolution, which can be crucial for characterizing cellular heterogeneity in the spatial context when investigating tissue organizations Miller *et al*. (2021); Moses and Pachter (2022). To address this limitation, spatial transcriptomics (**ST**) technologies can measure gene expression at a variety of spatial locations (spots) in a tissue sample while preserving the source position of each expression datapoint Marx (2021). Since the processes by which cells evolve into tissue compartments and interact with each other depend on interactions with the environment around it, spatial information which is naturally preserved by **ST** technologies presents ample opportunities for enhancing our understanding of disease progression and tissue development Walker *et al*. (2022).

Despite the rapid development of **ST** technologies, many of them still lack single-cell resolutions, such as Visium 10x Genomics (2022), Slide-seq Rodriques *et al*. (2019) and **HDST** Vickovic *et al*. (2019). In these approaches, each tissue is divided into a grid or lattice of spots, with each spot in the grid typically being 50–100*μ*m wide, covering around 10–60 cells. These **ST** technologies output a high-dimensional, spatially-localized gene expression count vector for each spot, representing an aggregated gene expression of the cells in the spot Zhang *et al*. (2023). As a result of the accumulated measurement at each detected spot, the measured signal is generally a mixture of multiple homogeneous or heterogeneous cell types, which may make it difficult to explore the spatial distribution of cell types in complex tissues Tu *et al*. (2023). Spot deconvolution methods aim to separate the contribution of different cell types in each spot, allowing for cell type identification and characterization. This enables the analysis of cell-type specific gene expression patterns and functional annotations, which is necessary for understanding the heterogeneity and cellular composition of complex tissues Ma and Zhou (2022). As a result of crucial need for methods capable of deconvolving cell type fractions for each spot to improve interpretability and analysis of gene expression patterns, recently several spot deconvolution tools have been developed such as **CARD** Ma and Zhou (2022), Bayes**TME** Zhang *et al*. (2023), **ST**deconvolve Miller *et al*. (2022), Cell2location Kleshchevnikov *et al*. (2022), Dest**VI** Lopez *et al*. (2022), **RCTD** Cable *et al*. (2022), **E**n**D**econ Tu *et al*. (2023), **SPOT**light Elosua-Bayes *et al*. (2021), and **U**ni**C**ell Charytonowicz *et al*. (2023).

One of the limitations of many existing spot deconvolution methods is the requirement for a reference profile of cell-type expression. Previous studies of **RNA**-seq data deconvolution algorithms have shown that choice of reference is more important than methods of choice in determining deconvolution performance. A reference-free spot deconvolution pipeline that does not rely on pre-existing reference atlases or datasets, assures an unbiased analysis of spatial transcriptomics data Cobos *et al*. (2020). **R**ecently, two reference-free tools, **ST**deconvolve and **B**ayes**TME** have been developed to deconvolve underlying cell types comprising multi-cellular spot resolution **ST** datasets Miller *et al*. (2022); Zhang *et al*. (2023). **ST**deconvolve is based on latent **D**irichlet allocation (**LDA**), a generative statistical model commonly used in natural language processing for discovering latent topics in collections of documents Miller *et al*. (2022). On the other hand, **B**ayes**TME** is a **B**ayesian hierarchical generative model capable of performing spot deconvolution for aggregated gene expression measurements at spots in **ST** datasets, explicitly modeling the aggregated counts via a Bayesian factorized model formulation Zhang *et al*. (2023).

While many of these **ST** analysis methods focus on analyzing individual **ST** samples, recent advances in high-throughput sequencing technologies, coupled with spatially resolved experimental techniques, have facilitated the generation of multi-sample **ST** datasets, enabling data integration and statistical modeling for more robust comparisons, validation, and identification of spatially regulated gene expression patterns Andersson *et al*. (2021); Mantri *et al*. (2021); Maynard *et al*. (2021a). For example, multi-sample **ST** allows more comprehensive investigation of gene expression spatial dynamics across different conditions (e.g., knock-out vs. wild-type) or experimental settings (e.g. treatment responders vs. non-responders) Allen *et al*. (2022). Additionally, Comparative analysis between samples offers insights into the spatial regulation of gene expression, unveiling spatial clusters and coordinated gene modules that would be overlooked in single-sample **ST** analysis. However, despite the ample opportunities that multi-sample **ST** data analysis may offer, to the best of our knowledge, there are no available spot deconvolution tools for integrative analysis of multi-sample **ST** datasets. **R**ecently, a hybrid machine learning and **B**ayesian statistical modeling framework called **MAPLE** has been developed for spot clustering of multi-sample **ST** data but does not perform spot cell-type deconvolution which is crucial for characterization of tissue samples Allen *et al*. (2022).

To fill these gaps, here, we introduce **MUSTANG**, a multi-sample spatial transcriptomics data analysis framework, to simultaneously derive the spot cellular deconvolution of multiple tissue samples without the need for reference cell type expression profiles. **MUSTANG** is designed based on the assumption that the same or similar cell types exhibit consistent gene expression profiles across samples. It adjusts for potential batch effects as crucial multi-sample experiments considerations to enable cross-sample transcriptional information sharing to aid in parameter estimation. With that, spatial correlation in gene expression patterns within samples is further accomodated by constructing and employing a spot “*similarity*” graph that includes both transcriptional and spatial similarity edges between spots across samples. By aligning and integrating multiple tissue samples, **MUSTANG** can effectively leverage shared information and increase the robustness of joint spot cell-type deconvolution analysis across multiple **ST** samples. In summary, our key technical contributions include:

- **MUSTANG** is the first reference-free spot deconvolution method for multi-sample **ST** data analysis, to the best of our knowledge.
- **MUSTANG** allows both intra-sample and inter-sample information sharing by introducing a new spot similarity graph.
- Besides modeling spot spatial dependency, MU**ST**ANT implements batch correction across **ST** samples in the workflow to avoid obscuring inherent biological signal when sharing transcriptional information.

To demonstrate the capability of **MUSTANG** for revealing the true underlying spot-level cell-type proportions in multi-sample **ST** datasets, we have applied **MUSTANG** to two real-world **ST** datasets of different tissue types and show that it can be effectively used for unveiling the inherent biological signal in tissue architectures.

## 2 Methods

Given gene count matrices of all spots across tissue samples and spatial coordinates for spot centroid positions, **MUSTANG** performs spot cellular deconvolution for multi-sample **ST** data. The overall workflow of **MUSTANG** is presented in Figure 1. **MUSTANG** includes four main steps: (i) construction of spot transcriptional adjacency matrix of expression-based information sharing across tissue samples after batch effect correction; (ii) construction of spot spatial adjacency matrix to allow spatial correlation between physically neighboring spots within the samples; (iii) construction of the spot “*similarity*” graph based on the spot transcriptional and spatial adjacency matrices; (iv) deconvolution of aggregated spot-level gene expression measurements to signals coming from different cell-types based on a Bayesian hierarchical model. Here, we discuss each step in more details, respectively:

**Figure 1:**
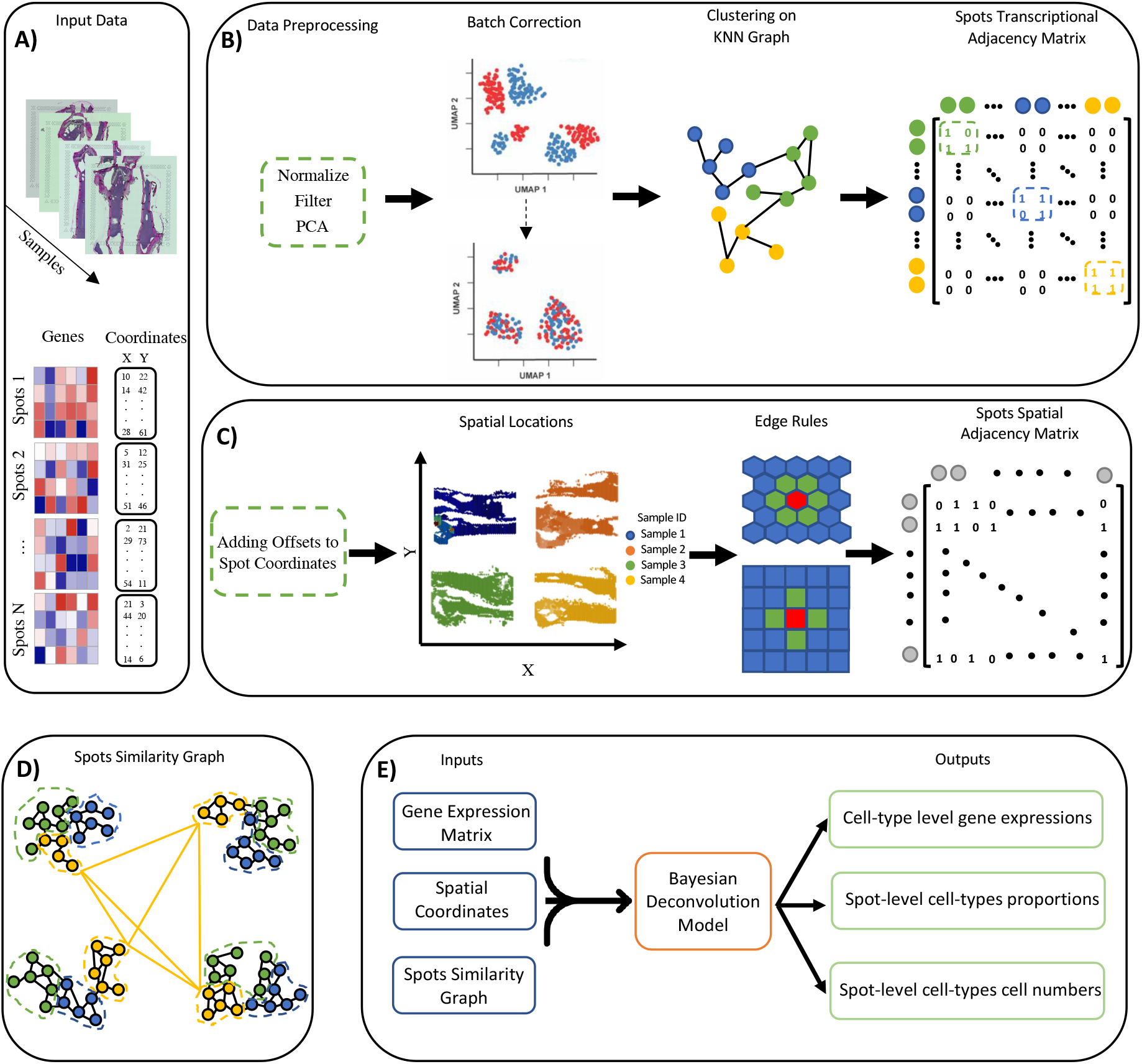
The **MUSTANG** framework to analyze multi-sample spatial expression data. A) **MUSTANG** requires gene expression matrices of all the spots across tissue samples as well as the spatial coordinates of the spots. The gene expression matrices are concatenated to form a single expression matrix of genes for all spots. B) **MUSTANG** performs standard sc**RNA**-seq data preprocessing steps such as normalization, gene filtering and then dimension reduction of gene expression matrices of the combined spots across samples via principal component analysis (**PCA**). The top principal components are batch-corrected to remove any unwanted technical confounders. Then **MUSTANG** performs Louvain clustering on the K-nearest neighbor graph constructed based on the batch corrected top **PC**s, to get the clusters of similar spots. The spot transcriptional adjacency matrix is then constructed based on the resulted spot cluster memberships. C) **MUSTANG** adds different offset values to the spatial coordinates of the spots from different **ST** samples so that they can be aligned properly. Depending on the sequencing technology layout (e.g. lattice or hexagonal), the spots spatial adjacency matrix is determined. D) The spot similarity graph is constructed by **MUSTANG** based on the summation of spots spatial and transcriptional adjacency matrices. Spots are colored by their corresponding transcriptional clusters. The edges in black indicate the spatial neighbouring connection between two spots and the yellow colored edges demonstrate the transcriptional similarity between yellow colored spots. E) Final step of **MUSTANG** corresponds to joint Bayesian deconvolution analysis based on raw concatenated gene expression matrix, spatial coordinates with added offsets, and the spot similarity graph.

### 2.1 Spot Transcriptional Adjacency Matrix

**MUSTANG** first identifies the common genes across multiple input tissue samples and then concatenates the spot count matrices of all samples *{*1, …, *N}* over the common genes (Figure 1.A). **T**hen, **MUSTANG** performs the common data preprocessing steps similar as typical sc**RNA**-seq data analysis, such as normalization, feature selection and dimension reduction. First, the combined gene expression matrix of all tissue samples are log transformed and normalized using library size. Then, the top 2000 (optional) highly variable genes are selected based on the variance of the log-expression profiles. We further perform **P**rincipal Component Analysis (**PCA**) on the normalized expression profiles of selected top highly variable genes across all the spots from tissue samples. Then, the reduced-dimension transcriptional matrix of all spots by top 50 principal components (**PC**) is retained to capture as much variation as possible while scaling up with complexity of analyzing high-dimensional data. In order to remove any unwanted technical batch effect from the analysis such as the case that tissue samples are from different sequencing technologies or samples are generated from multiple experiments or across different laboratories, **MUSTANG** performs batch effect correction on the retained top **PC**s. **O**ne powerful method for batch correction is the **H**armony algorithm Korsunsky *et al*. (2019). **MUSTANG** uses Harmony to adjust for batch effects from the **PC**s and ensures that the subsequent analyses are not confounded by technical variability. **L**ater, based on the batch corrected top 50 **PC**s, the **K**-nearest neighbor (**KNN**) graph of spots is constructed. Basically, in the **KNN** graph the nodes represent spots across **ST** samples and two spots are connected with an edge if they are within the k-most transcriptionally similar spots from each other for user-selected resolution parameter k. We measure the transcriptional similarity between spots by calculating the Euclidean distance of the batch corrected top 50 **PC** scores. Here, in **MUSTANG** we suggest selecting k to be 50 considering computation-performance trade-off. Additionally, we weigh the edges between two spots *i* and *j* in the **KNN** graph by 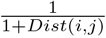where *Dist*(*i, j*) is the corresponding **PC**-based Euclidean distance between the two spots. This way, the edges between spots that are transcriptionally more similar will be weighed with higher values. Then, **MUSTANG** applies unsupervised graph-based Louvain clustering on the weighted **KNN** graph to get clusters of spots that are transcriptionally similar Blondel *et al*. (2008). Lastly, **MUSTANG** constructs the spot transcriptional adjacency matrix based on the spot membership in the resulted Louvain clustering results. If **T** is the cross-sample spot transcriptional adjacency matrix, then the value **T**_**ij**_ = **T**_**ji**_ = 1 at spots *i* and *j* means that *i* and *j* are in same transcriptional Louvain clustering class of spots and they are not within a same tissue sample (Figure 1.B).

### 2.2 Spot Spatial Adjacency Matrix

The next step in **MUSTANG** constructs spot spatial adjacency matrix. In this step **MUSTANG** only uses the coordinates of all the spots. Initially, we add different constant values to all spot coordinates of different samples so that it could be possible to overlay the physical locations of spots from different samples on a single layout without spots from different samples getting overlapped or neighboured as shown in Figure 1.C. Then, based on the geometric representations of spots in **ST** sequencing technologies, such as lattice layouts (e.g., Slide-seq Rodriques *et al*. (2019)) or hexagonal layouts (e.g., Visium 10x Genomics (2022)), neighbors can be identified for each spot based on shared edges. This edge rule leads to four and six neighbors for non-boundary spots in lattice and hexagonal layouts, respectively. Finally, **MUSTANG** constructs the spots spatial adjacency matrix based on the described edge rule. If we call the spots spatial adjacency matrix **S**, then the value **S**_**ij**_ = **S**_**ji**_ = 1 means that *i* and *j* have a shared edge between them (Figure 1.C).

### 2.3 Spot Similarity Graph

After deriving both spot transcriptional and spatial adjacency matrices, **MUSTANG** constructs the overall spot “*similarity*” graph. The adjacency matrix of spot similarity graph is a binary matrix which is resulted after taking the logic “OR” operation between pairwise indices of spot transcriptional and spatial matrices **T** and **S**. More specifically, if we denote the spot similarity graph adjacency matrix by **A, A**_**ij**_ = **T**_**ij**_*∨* **S**_**ij**_, where “*∨*” indicates the “OR” operator. Figure 1.D shows an example of how a spot similarity graph might look like for a **ST** dataset with four tissue samples. In this figure, spots are colored based on their transcriptional cluster labels. The black colored edges are the edges according to the spot spatial adjacency matrix. On the other hand, the yellow colored edges indicate the transcriptional similarity between yellow colored spots. Note that for simplicity, only the transcriptional edges between yellow colored spots are drawn and transcriptional edges between blue and green spots are not shown in the figure. Additionally, it worth mentioning that each yellow edge between a pair of yellow spots in the corresponding clusters is representative of all edges from spots of one cluster to another in Figure 1.D

### 2.4 Joint Bayesian Deconvolution Analysis

The last step of our **MUSTANG** workflow corresponds to joint Bayesian deconvolution analysis of raw concatenated gene expression matrix to preserve information in the original **ST** data, together with the spot similarity graph and spatial coordinates with added offsets. Our joint Bayesian deconvolution model, is based on the Poisson discrete deconvolution model recently introduced in **B**ayes**TME** for single sample analysis of **ST** data Zhang *et al*. (2023). More precisely, in this Poisson model, the raw aggregated expression measurement of gene *g* at spot *s*, denoted as *Y*_*sg*_, are factorized as the summation of *k* (i.e. number of cell types) different Poisson distributed read counts *Y*_*sgk*_. In fact, each of these reads models the total expression count of gene *g* in the cells of type *k* that are at spot *s*. Thus, based on this factorization we can explicitly model the raw **ST** counts *Y*_*sg*_:

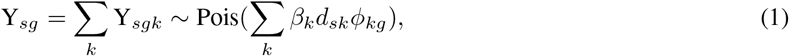

where the rate parameter of the Poisson distributions is controlled with three parameters *β*_*k*_, *d*_*sk*_ and *ϕ*_*kg*_. The cell type dependent parameter *β*_*k*_ quantifies the expected total count for cells of type *k* and *d*_*sk*_ represents the number of cells of type *k* that are at spot *s*. The parameter *ϕ*_*kg*_ captures the normalized gene expression profile of gene *g* in cell type *k*. This way of modeling gene expression in **ST** data assures biological considerations such as monotonic relationship between the number of cells and aggregated read measurement in each spot as well as different expression profiles for each gene in various cell types. To complete the Poisson discrete deconvolution model, Dirichlet and gamma distribution priors are imposed on *ϕ*_*k*_ and *β*_*k*_ parameters, respectively. Additionally, the prior on *d*_*sk*_ is constructed hierarchically based on the heavy-tailed **B**ayesian variant of the graph-fused **B**inomial tree as described in Tansey *et al*. (2017). In this Binomial tree model, the cell type assignment probabilities in each spot are decomposed into a series of binomial decisions where the prior on each binomial probability encourages spatial smoothness across spots. Specifically, such spatial smoothness on cell type assignment probabilities is achieved by imposing the sparsity inducing grouped horseshoe distribution Xu *et al*. (2016) over the graph fussed **LASSO** Wang *et al*. (2016) (i.e. zeroth-order graph trend filtering) penalized cell type assignment probabilities:

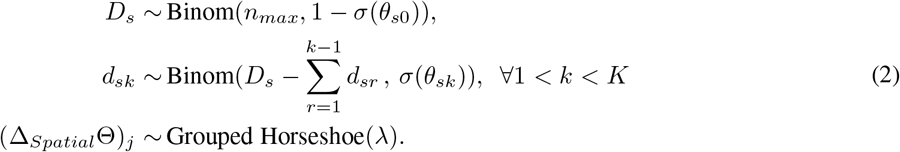

In equation (2), *n*_*max*_ is the maximum possible number of cells in each spot, for which we set to be 100 in our experiments. The parameter *D*_*s*_ is the total number of cells in spot *s* out of possible *n*_*max*_ cells and *θ*_*sk*_ captures the cell type *k* probability proportions at spot *s*. Lastly, Δ_*Spatial*_ is the edge-oriented zeroth-order graph trend filtering matrix of the spot spatial graph with a hyper-parameter *λ* controlling the global degree of smoothness.

**H**ere, in our joint **B**ayesian deconvolution model while performing multi-sample **ST** data analysis in **MUSTANG**, we further allow information sharing across tissue samples in the **P**oisson discrete deconvolution model. We take advantage of the prior knowledge inherited in the spot similarity graph that we constructed in the **MUSTANG** workflow as detailed in the previous section. Specifically, we include transcriptional similarity in addition to the spatial similarity to take into consideration of the biological belief that spots that have similar batch corrected transcriptional profiles might also have similar cell type composition as well. This is done by taking advantage of the zeroth-order graph trend filtering matrix of the spot similarity graph in the hierarchical prior in equation (2). In **MUSTANG**, we impose the grouped horseshoe distribution over the graph fussed **LASSO** penalized cell type assignment probabilities based on the spot similarity graph as:

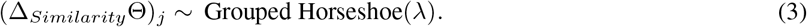

This results in inferring both transcriptionally and spatially smooth cell type proportions, allowing to borrow signal strengths from both inter-sample and intra-sample spots for effective joint analysis of multiple tissue samples in a given **ST** dataset.

The posterior inference procedure of the joint Bayesian deconvolution model in **MUSTANG** is based on Gibbs sampling. The full derivations for all complete conditionals and Gibbs sampling based updates are similar as Zhang *et al*. (2023) and detailed in the supplementary materials. During the inference process, we use **M**arkov chain thinning with five thinning steps between each sample. **W**e collect 100 **M**arkov chain **M**onte Carlo (**MCMC**) samples after 1200 burn-in iterations for our consequent analyses and evaluation.

## 3 Results & Discussion

We have evaluated our **MUSTANG** for analysis of multi-sample **ST** data from two real-world **ST** datasets generated by the 10X Genomics Visium platform 10x Genomics (2022). First, a human brain **ST** dataset is used to quantitatively benchmark the spot deconvolution performance of **MUSTANG**. Specifically, the significance of different components in **MUSTANG** enabling multi-sample **ST** analysis will be demonstrated in this ablation study comparing with Bayes**TME** and a simpler version of **MUSTANG** that does not take spot transcriptional adjacency matrix into account. We then analyze a mouse bone marrow tissue **ST** dataset to characterize tumor microenvironment. The matched immunofluorescence (**IF**) staining images are used to validate the findings by analyzing bone tissue samples with **MUSTANG**.

### 3.1 Human Brain Data

In a recent study Maynard *et al*. (2021b), spatial expression profiles of 12 dorsolateral prefrontal cortex (**DLPFC**) tissue samples were generated. Based on the selected **DLPFC** layer-specific gene makers and cytoarchitecture consideration, six cortical layers (i.e. **L1-L6**) and white matter (**WM**) for each brain tissue sample were annotated. Here, we use the **ST** expression profiles of four samples (Sample **ID**: 151673 to 151676) from this dataset to showcase the benefits of simultaneously denconvolving tissue samples using our proposed **MUSTANG**.

Figure 2.A shows the Hematoxylin and Eosin (**H&E**) staining images of four **DLPFC** tissue samples from the human brain **ST** dataset as well as the cortical layers and white matter reference annotations for sample 151673 from the original study. Following our **MUSTANG** workflow, we first start analyzing the samples by constructing spot transcriptional and spatial adjacency matrices. As shown in Figure 2.B, we derive the spot spatial adjacency matrix by adding offsets to spatial coordinates of **DLPFC** tissue samples and overlaying them on the **ST** grid space based on the Visium platform. In the transcriptional space, we follow the data preprocessing steps previously described in Section 2.1 to derive the dimension-reduced top 50 **PC**s for spot-aggregated gene expression counts. Figure 2.C displays the **UMAP** (Uniform Manifold Approximation and Projection McInnes *et al*. (2020)) embedding of the derived top 50 **PC**s. It can seen that there is strong batch effect in this dataset as spots from different tissue samples are clustered based on the their sample **ID** rather than their underlying biological cell types. Although these samples are from the same tissue and sequencing platform, this observed batch effect in the data calls for the need of batch effect correction when analyzing multiple tissue samples to reduce the potential influence from any confounding technical factor. We therefore implement Harmony in **MUSTANG** to derive the batch corrected top 50 **PC**s. The **UMAP** embedding of the batch corrected **PC**s are shown in Figure 2.D, where the spots from different samples are now mixed together while preserving potential expression differences. We further construct the **KNN** graph of spots based on these top **PC**s and apply Louvain clustering, resulting 8 distinct transcriptional sub-populations. In Figure 2.E, the spots from four samples are colored by their transcriptional clusters in the **UMAP** embedding space. With that, the spot transcriptional adjacency matrix and consequently, the spot similarity graph, can be constructed. Finally, we fit our joint Bayesian deconvolution model to the concatenated data with *K* = 7 cell types (i.e. six cortical layers plus white matter). Based on the collected post burn-in **MCMC** samples, we derive the posteriors of the joint deconvolution model parameters such as spot-wise cell type proportions, cell types cell numbers and normalized cell-type specific gene expression. Figure 2.F demonstrates the spatial scatter pie chart plot of our four **DLPFC** tissue samples, in which spots are plotted in their physical coordinates and at each spot there is a circular pie chart representing the inferred proportions of assigned cell types in that spot. The high similarity between the spatial patterns of cell type proportions in the spatial pie chart plots of all four samples and the ground truth annotations from the original study demonstrates the capability of **MUSTANG** to simultaneously infer the underlying spot-wise biological cell type proportions across multiple tissue samples.

**Figure 2:**
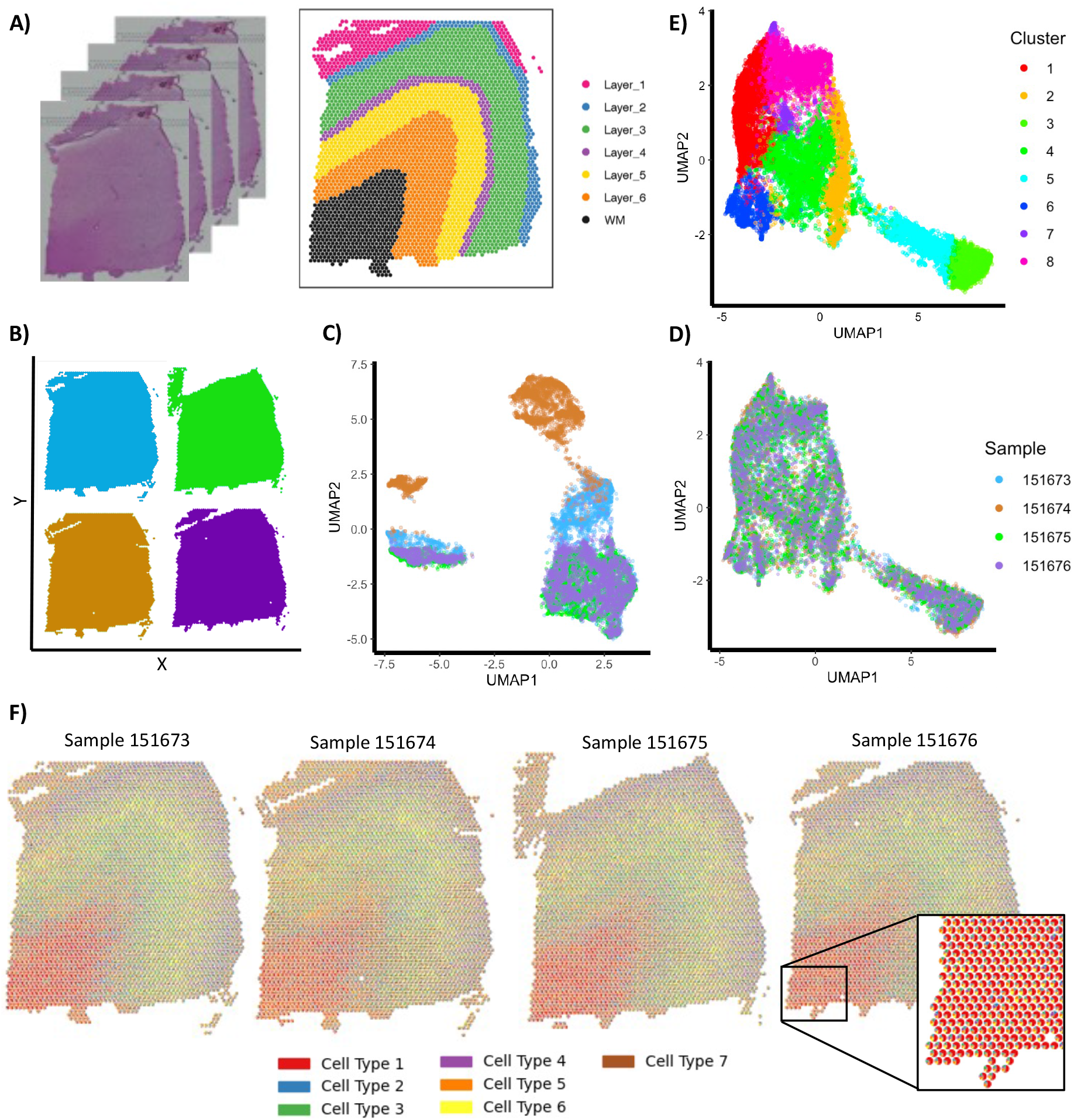
Analysis of four human brain **DLPFC** tissue samples with **MUSTANG. A) H&E** staining images of four tissue samples (right) and the reference annotations of spots for the sample 151673 (left). **B)** Overlaying tissue samples on a grid space to construct spot spatial adjacency matrix. **C) UMAP** embedding visualization of spots by top 50 **PC**s before and **D)** after batch correction. **E)** Visualization of clustering based on batch corrected top 50 **PC**s. **T**he spots are colored based on their transcriptional cluster label inferred from Louvain clustering. **F)** Spot-based spatial pie charts of **MUSTANG**-inferred cell type proportions across all four **DLPFC** tissue samples matching the reference annotations from the original study.

As the ground truth cell type proportions and cell type cell numbers does not exist for multi-cell resolution **ST** data, inspired by the guidelines described in the recent benchmarking study of cell type deconvolution methods for **ST** data Li *et al*. (2023), we quantify the cell type cell number inference performance of **MUSTANG** based on the Pearson correlation coefficient (**PCC**) between the predicted spot-wise cell counts of specific cell type (i.e. *d*_*sk*_ in Equation 1) and the corresponding marker genes’ expression profiles. Specifically, we benchmark **MUSTANG** with **B**ayes**TME**, which is a **ST** data deconvolution tool capable of inferring cell type cell numbers without the need for paired reference expression profiles. As **B**ayes**TME** is designed for single sample analysis, we analyze each brain tissue sample separately using **B**ayes**TME** as the baseline.

To calculate the **PC**C values, we first gather the list of known layer-specific marker genes from two previous brain studies Molyneaux *et al*. (2008); Zeng *et al*. (2012) that were also used in the original **DLPFC** dataset paper Maynard *et al*. (2021b). Specifically, we only use those marker genes that are annotated to be related to only one of the **DLPFC** layers except for white matter (**WM**) layer, for which as we could not find any **WM-**specific markers in the two references, we select the marker genes that are shared between layer 6 and **WM**. The heatmap plot in Figure 3.A shows the list of selected layer-specific marker genes. The colors in the plot represent the corresponding reference papers that reported the corresponding marker genes.

**Figure 3:**
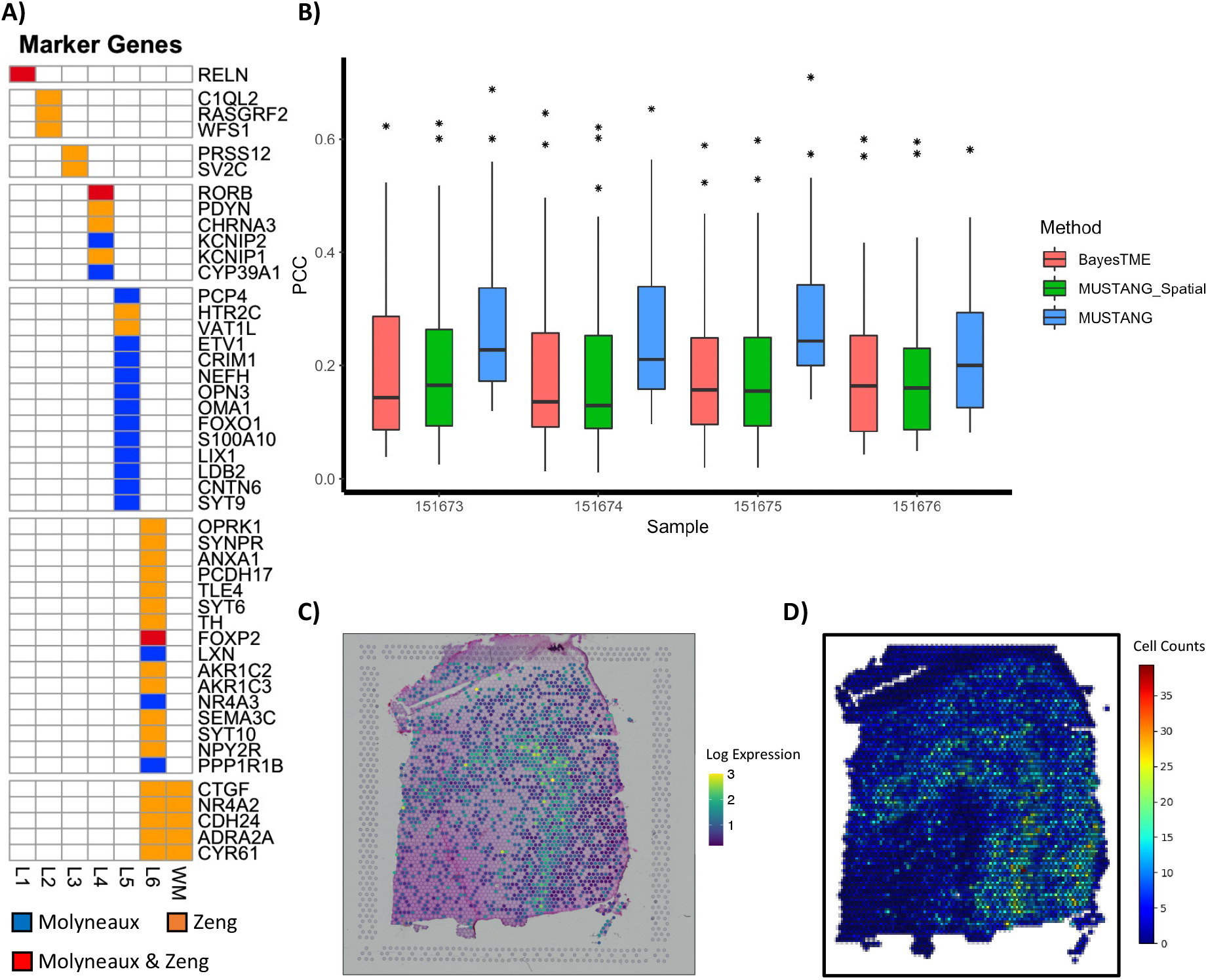
Quantitative performance benchmarking on four **DLPFC** tissue samples. **A)** List of layer-specific gene markers from two brain tissue studies Molyneaux *et al*. (2008); Zeng *et al*. (2012). **B)** Boxplots showing the calculated **PC**C values for three different reference-free cell type deconvolution methods: **MUSTANG, MUSTANG**_Spatial and **B**ayes**TME**. Higher **PC**C values indicate better deconvolution performance identifying annotated cell types. **C)** Spot-level log2 expression visualization of the **L5** layer marker gene **PCP4** correlates with the spatial pattern of **D) MUSTANG-**inferred cell numbers for the **L5** layer best paired cell type for the sample 151674.

Next, we extract the layer-specific gene expression profiles of **DLPFC** layers based on the “*pseudo-bulking*” approach noted in the original study of the **DLPFC** dataset Maynard *et al*. (2021b), in which the **UMI** counts for each gene within each layer across 12 spatial replicates are summed up to generate layer-enriched expression profiles. The layer-specific gene expression profiles of **DLPFC** layers have shown previously in Maynard *et al*. (2021b) to capture biological properties inherent in **DLPFC** layers. This pseudo-bulk data is availabe as “sce_layer data” for download through the `fetch_data` function in `spatialLIBD` R package. Following the instructions for cell type deconvolution benchmarking described in Li *et al*. (2023), for each **DLPFC** layer, we calculate the **PC**C between the expression profile of each layer in the extracted pseudo-bulk data and the inferred normalized expression profile of all cell types (i.e. *ϕ*_*kg*_ in Equation 1) from **MUSTANG**, choose the best-paired inferred cell type with the highest **PC**C and match it to that layer. After assignment, this chosen cell type would be ignored in the future steps. Then, we repeat the aforementioned steps on the next layer until all layers are iterated. For now, each layer should be paired with the best suitable cell type without duplication.

Finally, to complete the quantitative comparison between different **ST** analysis methods, for each **DLPFC** layer, we calculate the **PC**C value between the corresponding marker gene expression of that layer in Figure 3.A and the inferred expression corresponding to the best-paired cell type. we calculate **PC**C values for each of the four tissue samples separately after jointly analyzing them with **MUSTANG**. We repeat the same procedure for analyzing tissue samples separately using **B**ayes**TME** and calculate the corresponding **PC**C values. The boxplots in Figure 3.B shows the **PC**C values for each method on each sample separately. As depicted in the figure, on all four tissue samples, jointly analyzing them with **MUSTANG** leads to higher average **PC**C values comparing to separately deconvolving them using **B**ayes**TME**. This superior performance of **MUSTANG**, illustrates the benefit of simultaneously analyzing tissue samples with an approach that allows for effective cross-sample information sharing. As an example of the spatial expression pattern of the marker genes and inferred cell-type cell numbers, we have visualized the log2 expression of the ***L5*** layer marker gene ***PCP4*** as well as the **MUSTANG** inferred cell numbers for the **L5** layer best paired cell type for the sample 151674 in Figures 3.C and 3.D, respectively. The derived **PC**C value for this gene is 0.42. **H**ere, we would like to emphasize that due to the nature of quantitative analysis we did in this section while **ST**deconvolve deconvolution model does not explicitly model cell type cell numbers (i.e. *d*_*sk*_ in our deconvolution model), it is not possible to benchmark **ST**dencovolve with other comparing methods for the presented performance evaluation results. It worth mentioning that adjusting for this parameter during the deconvolution of aggregated **ST** signals in multi-cellular spot resolution **ST** datasets is crucial to assure biological considerations such as monotonic relationship between the number of cells and aggregated read measurement in each spot. As currently, to the best of our knowledge only **MUSTANG** and **B**ayes**TME** adjust for this source of variation, we have only included these methods results in Figure 3.B and excluded **ST**deconvolve from this quantitative analysis.

To better understand the corresponding contributions of different components in **MUSTANG** to its superior performance for multi-sample **ST** data analysis, we further conduct the ablation study that analyzes the tissue samples with a simplified version of **MUSTANG** without using the spot transcriptional adjacency matrix across samples. This means that we deconvolve tissue samples without cross-sample transcriptional information sharing. We call this simpler version of **MUSTANG**, “**MUSTANG**_**S**patial” as after removing transcriptional edges from spot similarity graph, it gets reduced to using only the spot spatial coordinates. As shown in Figure 3.B, the **PC**C values in all four samples get significantly lower in the obtained results by “**MUSTANG**_Spatial” in comparison with those by the complete **MUSTANG** workflow. **C**learly, removing transcriptional information sharing from **MUSTANG**, leads to on average similar **PC**C values of the results using **B**ayes**TME** that deconvolves tissue samples separately. This is expected as **B**ayes**TME**, similar as “**MUSTANG**_Spatial”, only allows within-sample information sharing across physically neighboring spots by performing spatial smoothing on cell type assignment probabilities. **T**his ablation study, clarifies the significance of intra-sample transcriptional similarity guidance on boosting the performance of **MUSTANG**.

### 3.2 Mouse Bone Marrow Data

The tumor microenvironment (**TME**) plays a critical role in tumor development, progression, and therapeutic response Kalbasi and Ribas (2020). **R**ecently, several studies have reported that the spatial organization of **TME** is the key determinant of the disease behavior and treatment outcomes Fu *et al*. (2021); Blise *et al*. (2022). **T**hus, a comprehensive understanding of the spatial architecture and expression patterns of **TME** holds great promise for the development of novel therapeutic treatment strategies. **T**aking advantage of the **TME** spatial transcriptomics data helps unveil the underlying complex spatial organization and intricate interplay between tumor cells and their microenvironment.

For the second application of **MUSTANG** analyzing **ST** data of tissue samples, we study and characterize mouse bone marrow tissue tumor microenvironment. We have profiled the bone tissue of 6-8 weeks mouse after bone lesions generation by Intra-iliac injection (**IIA**). For spatial analysis, **ST** data are obtained via the 10**X** Visium platform to profile four bone marrow tissue sections. Specifically, thin (10-*μ*m) mouse bone marrow sections were mounted directly onto separate designated capture areas on the 10x Visium spatial gene expression slides and data preprocessing was done per the manufacturer’s protocols. In brief, after **H&E** staining, each section was imaged using color bright field by Cytation 5. The sections were then processed following the 10X Visium gene expression protocols until the c**DNA** libraries were constructed, which were later sequenced by Novaseq 6000 system with 150bp paired end reads, aiming at 300 million raw reads per section. The **H&E** staining images of the four bone tissue sections are shown in Figure 4. The Visium **S**patial **G**ene **E**xpression **S**olution from 10x Genomics allows for the analysis of m**RNA** using high-throughput sequencing and subsequently maps a transcript’s expression pattern in tissue sections using high-resolution microscope imaging. **T**his provides gene expression data at 5,000 capture spots in each Visium slide within the context of tissue architecture, tissue microenvironments, and cell groups. **S**pace**R**anger was used to process Visium spatial **RNA**-seq output and brightfield and fluorescence microscope images to detect tissue, align reads and generate feature-spot matrices. **S**pace**R**anger built-in function `mkfastq` was used to wrap Illumina’s `bcl2fastq` to correctly demultiplex Visium-prepared sequencing runs and to convert barcode and read data to **FASTQ** files. **S**pace**R**anger function `count` was used to take a microscope slide image and **FASTQ** files from **S**pace**R**anger `mkfastq` and perform alignment, tissue detection, fiducial detection, and barcode/**UMI** counting. In our study, raw sequence reads were mapped to mice reference genome (mm10) to obtain the gene expression profile at each spot.

**Figure 4:**
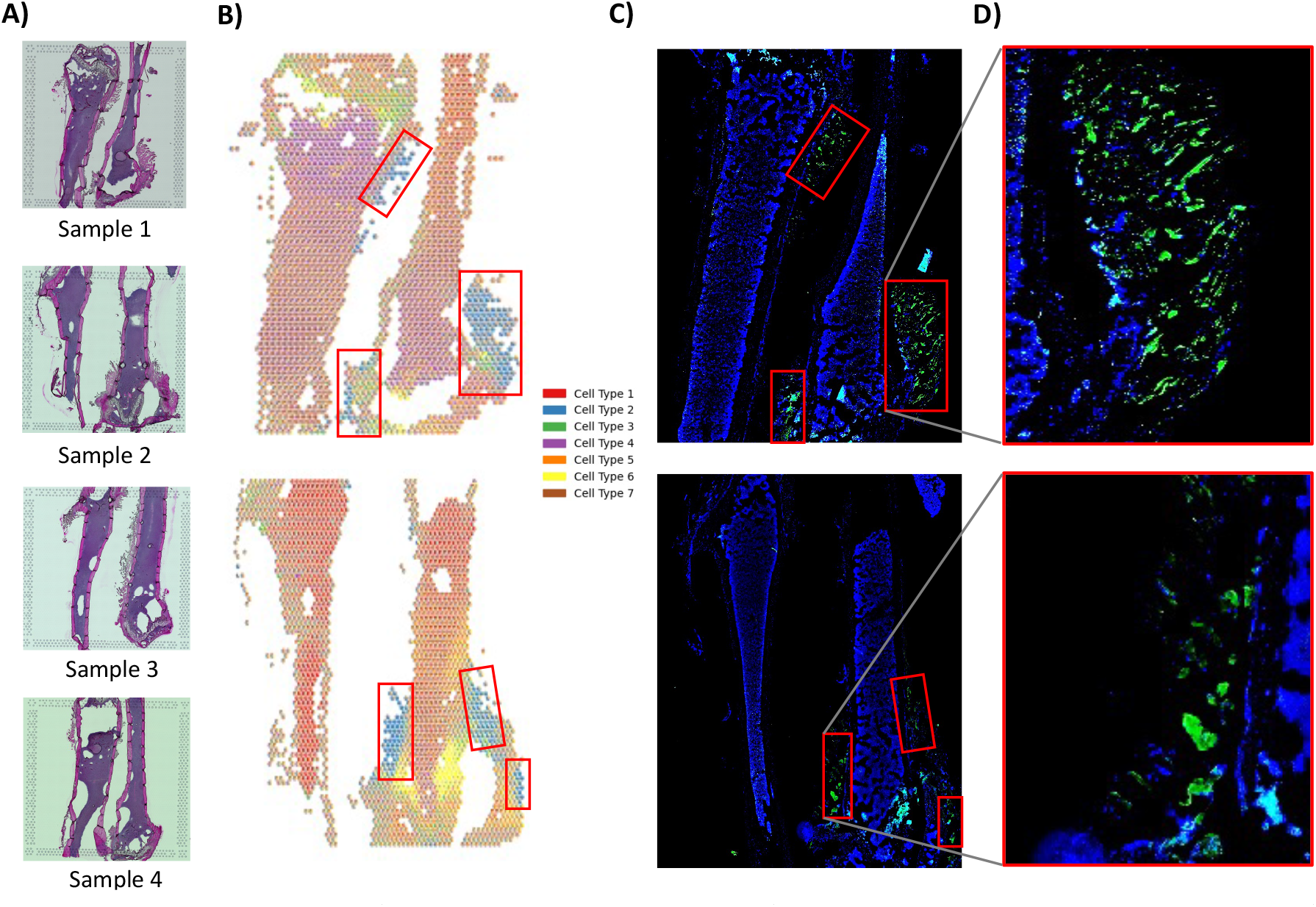
Analysis of four mouse bone marrow tissue samples with **MUSTANG. A) H&E** staining images of the four samples profiles with Visium platform. **B**) Spot-based spatial pie charts of **MUSTANG**-inferred cell type proportions for (top) sample 1 and (below) sample 2. **C) M**atching **IF** staining images of (top) sample 1 and (below) sample 2. **D)** Closer look at the IF staining image regions with high density of green dots, indicating the presence of tumor cells.

In order to identify and characterize the spatial organization of tumor cells within the bone marrow tissue **TME**, we jointly analyze the **ST** data from the four bone tissue sections with **MUSTANG. W**e follow the same **MUSTANG** workflow steps described in detail in section 2 to infer the deconvolved components of the bone tissue samples. We pick the number of cell types *K* based on the results of applying unsupervised cell type number inference algorithms implemented in **B**ayes**TME** Zhang *et al*. (2023) and **ST**deconvolve Miller *et al*. (2022) to each of the individual four bone tissue samples leading to 8 different inferred number of cell types. We then select the *K* to be 7 as it is the most frequently inferred value of total cell type numbers out of the eight derived values by **B**ayes**TME** and **ST**deconvolve. (Four occurrence; **D**etails in the supplementary materials).

After simultaneously analyzing the four bone tissue samples using **MUSTANG**, we plot the spatial scatter pie chart visualization of the inferred deconvolved cell type proportions. The spatial pie chart plots for samples one and two are visualized in Figure 4.B. To validate the identification of tumor cell types in the bone marrow **TME** by **MUSTANG**, we generate matched immunofluorescence (**IF**) staining images for each bone tissue sections separately. Specifically, the bone sections were stained with antibodies to depict the potential tumor cell enriched tissue section parts (The detailed protocol for generation of **IF** staining images can be found in the supplementary materials). The generated **IF** staining images for bone tissue samples one and two are shown in Figure 4.C. The green dots in the **IF** staining images highlight the tumor cell enriched parts (Figure 4.D). Matching the green dots regions in **IF** staining images with the spatial pie chart plots of tissue samples from **MUSTANG**, revealed the presence of high **MUSTANG**-inferred proportions of cell type 2 (colored with blue in Figure 4.B). We plot red boxes to highlight the regions of **IF** staining images of bone tissue samples with high enrichment of green dots (i.e. tumor cells) and overlay the boxes on the spatial pie charts. The spots in the matching red boxes of the spatial pie charts are composed of high inferred cell type number 2 proportions with **MUSTANG**. This demonstrates the capability of **MUSTANG** to identify tumor cell type cells in the bone marrow **TME**.

## 4 Conclusions

We have developed **MUSTANG**, a multi-sample **ST** data analysis workflow that jointly analyzes multiple tissue samples by leveraging transcriptional information sharing across samples as well as spatial dependency in gene expression patterns within samples.

By our proposed workflow, including spot similarity graph construction and batch effect correction removing unwanted nuisance factors obscuring the inherent biological signal in **ST** data, the joint **B**ayesian decovolution model in MUS-TANG extends the previous developments for reference-free single-sample **ST** data analysis Zhang *et al*. (2023) to joint multi-sample **ST** data analysis, allowing for robust simultaneous spatial characterization of cell sub-populations across spots in all tissue samples. **W**e have introduced a new spot-based knowledge graph, spot similarity graph, that captures sufficient and comprehensive similarity information between spots to be used in our joint **B**ayesian deconvolution model to improve the multi-sample analysis performance beyond existing methods analyzing single **ST** samples separately. **B**y providing extensive results on two real-world multi-sample **ST** data, we have demonstrated the superior performance of **MUSTANG** in terms of cell type deconvolution and spatial characterization of complex tissue environments. Future work concerns further improving the capability of **MUSTANG** to decipher tissue structures by performing joint cell-cell interaction analysis between cells of different sub-populations across multi-sample tissue samples.

## A. Supplementary Materials

### A.1 Gibbs sampling inference

Here, we provide the detailed posterior Gibbs sampling procedure for the joint **B**ayesian deconvolution model described in section 2.4.

#### Sampling *Y*_*sgk*_

Since we are modeling the raw **ST** counts *Y*_*sg*_ as

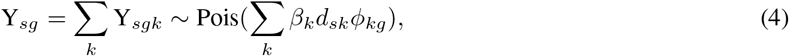

and leveraging the relationship between the Poisson and Multinomial distribution, the *Y*_*sgk*_ parameters can be sampled from a Multinomial distribution. If we define the auxiliary variables 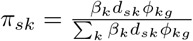, then

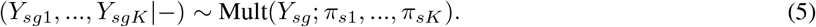

#### Sampling *β*_*k*_

To infer the cell type dependent expected total counts parameter *β*_*k*_, we write its posterior as

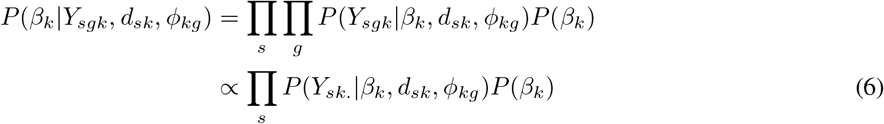

where *Y*_*sk*._ is (*Y*_*sk*1_, …, *Y*_*skG*_). Then, we can write the likelihood of reads *Y*_*sk*._ As

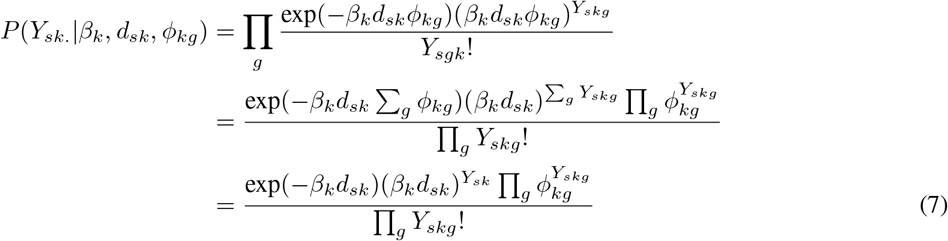

where in the last equation we take advantage of facts that Σ _*g*_ *ϕ*_*skg*_ =1 and Σ _*g*_ *Y*_*skg*_ *=Y*_*sk*_. Now, based on Equation 7, we can simplify the posterior of cell type dependent parameter *β*_*k*_ in Equation 6 as

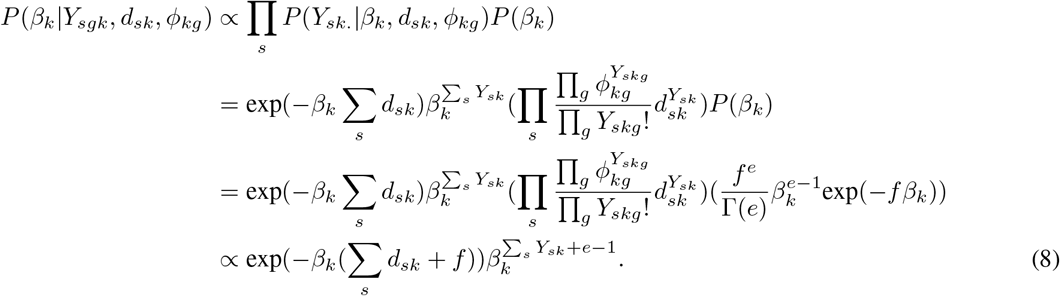

Note that in Equation 8, we leverage the Gamma prior distribution (i.e. Gamma(*e, f*)) we imposed on *β*_*k*_ as described in the main text. Thus, based on Equation 8 we can update the *β*_*k*_ as

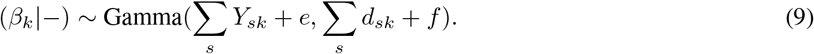

#### Sampling *ϕ*_*k*_

As described in the main text, we impose Dirichlet prior distribution over the normalized cell type dependent gene expression profile parameter *ϕ*_*k*_ = (*ϕ*_*k*1_, …, *ϕ*_*kG*_) (i.e. *ϕ*_*k*_ *∼* Dir(*α*_*k*_)) and _*g*_ *ϕ*_*kg*_ = 1. We have

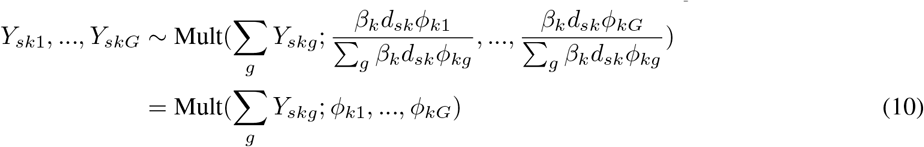

Thus, the normalized gene expression profiles can be updated using the Dirichlet-Multinomial conjugacy as

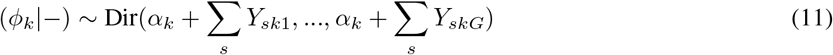

#### Sampling *D*_*s*_ and *d*_*sk*_

**B**y modeling the cell number distribution as hidden markov model (HMM) and exploiting the forward-filtering backward-sampling algorithm introduced in Zhang *et al*. (2023), we can update *d*_*sk*_ in efficient approach. Specifically, in the forward-filtering algorithm we calculate the “alpha” values of our hidden latent stats which includes the cell type cell numbers (i.e. *x*_*k*_) which we define as

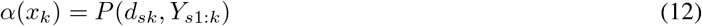

and in the backward-sampling, based on the derivations in Zhang *et al*. (2023), the cell type cell number values are updated based on

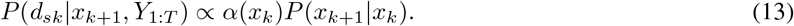

### A.2 Inferring total number of cell types (*K*) for mouse bone marrow data

Here, we describe the results of applying unsupervised cell type number inference algorithms implemented in **B**ayes**TME** Zhang *et al*. (2023) and **ST**deconvolve Miller *et al*. (2022) to each of the individual four mouse bone marrow tissue samples. **B**ased on the instructions in Miller *et al*. (2022), to find optimal number of cell types in bone tissue samples with **ST**deconvolve, we fit a number of different **L**atent **D**irichlet allocation (**LDA)** models with different **K** values and then based on the inferred number of “rare” predicted cell-types and perplexity values, we pick the number of cell types. Specifically, we change *K* from and 2 to 15 for each bone tissue sample and plot the perplexity and number of “rare” predicted cell types versus the *K* values. Figure S1 shows the **ST**deconvolve inferred perplexity and number of “rare” cell types versus different *K* values for four bone marrow samples one to four respectively. As described in **ST**deconvolve workflow Miller *et al*. (2022), we pick the number of cell types to be the value from that perplexity stabilizes and has the lowest number of rare sub predicted cell types to avoid over-clustering. **T**his leads to inferring 6, 7, 6 and 7 number of cell types for Samples 1 to 4 respectively (Table 1).

**Table 1:** Inferred total number of cell types (*K*) from **ST**deconvolve and **B**ayes**TME**.

| Method | Sample 1 | Sample 2 | Sample 3 | Sample 4 |
| --- | --- | --- | --- | --- |
| STdeconvolve | 6 | 7 | 6 | 7 |
| BayesTME | 8 | 7 | 7 | 8 |

Then, we use **B**ayes**TME** to infer total number of cell types (*K*). Specifically, **B**ayes**TME** does this by performing 5-fold cross-validation for each *K* = (2, …, 12) values with 5% of spots held out in each fold. Then, in each fold, a **P**oisson based discrete deconvolution model is fitted over a discrete grid of *λ* smoothness values (10^0^, 10^1^, …, 10^5^) and average log-likelihood for the held out spots are calculated. Finally, the *K* with highest averaged likelihood is picked to be the total number of cell typesZhang *et al*. (2023). Figure S2 shows the calculated average cross-validation log-likelihood versus the number of cell types for each of four bone tissue samples. **B**ased on these figures, the inferred total number of cell types for samples 1 to 4 is 8, 7, 7 and 8 respectively (Table 1).

Table 1 summarizes the inferred total number of cell types from **ST**deconvolve and **B**ayes**TME**. We then select the *K* to be 7 in our multi-sample analysis with **MUSTANG** as it is the most frequently inferred value of total cell type numbers out of the eight derived values.

### A.3 Immunofluorescence (IF) staining images generation protocol

Here, we describe the protocols for immunofluorescent staining of thick sections and bone clearing. **B**riefly, femur bone sections were cleaned, pretreated with 1mg/ml sodium borohydride solution and then blocked before whole-mount staining. Then, the bone sections were stained with antibodies. Immunofluorescent staining were performed within 1 ml staining buffer for three days at 4°C with constant rotation and followed by a whole day of PBS washing. The stained samples were then dehydrated by a series of methanol solutions before completely cleared by BABB solution. The bone sections were later sealed in imaging glass cassettes with BABB solution. The images were taken by an Olympus FV1200 MPE confocal microscope.

### A.4 Additional results with mouse bone marrow data

Here, we present the additional results of jointly analyzing four bone tissue samples as well as the **IF** staining images for the profiled tissue samples that highlights the tumor cells. Specifically, here, we focus on mouse bone marrow tissue samples 3 and 4 as the results of other two samples are discussed in detail in section 3.2. Figure S3.A shows the spatial pie chart plots generated by **MUSTANG** for samples three and four and sames as what we described in section 3.2, the **IF** staining images are generated and used to validate **MUSTANG** results by identifying tumor cells in bone marrow **TME**. Figure **??**.B shows the matched **IF** staining images for the bone marrow tissue samples three and four. As the figures suggest the green dots that highlight the tumor cells regions can be matched with the tissue areas in samples three and four that have high proportions of cells of cell type 2, illustrating the capability of **MUSTANG** to characterize tumor cells in mouse bone marrow **TME**.

---

**Figure S1:**
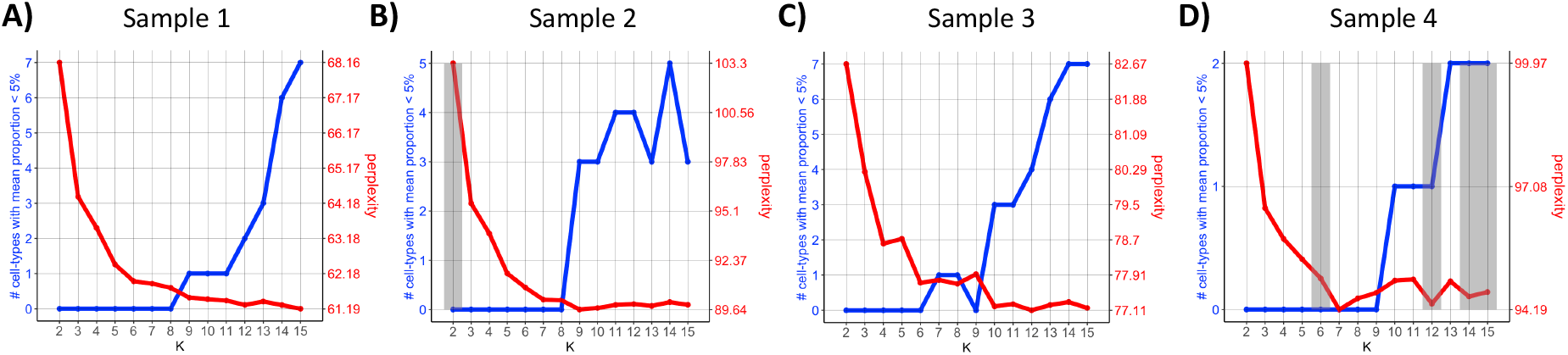
Inferring the number of cell types with **ST**deconvolve. Inferred perplexity and number of “rare” cell types are plotted versus different *K* values for four profiled bone marrow tissue samples.

**Figure S2:**
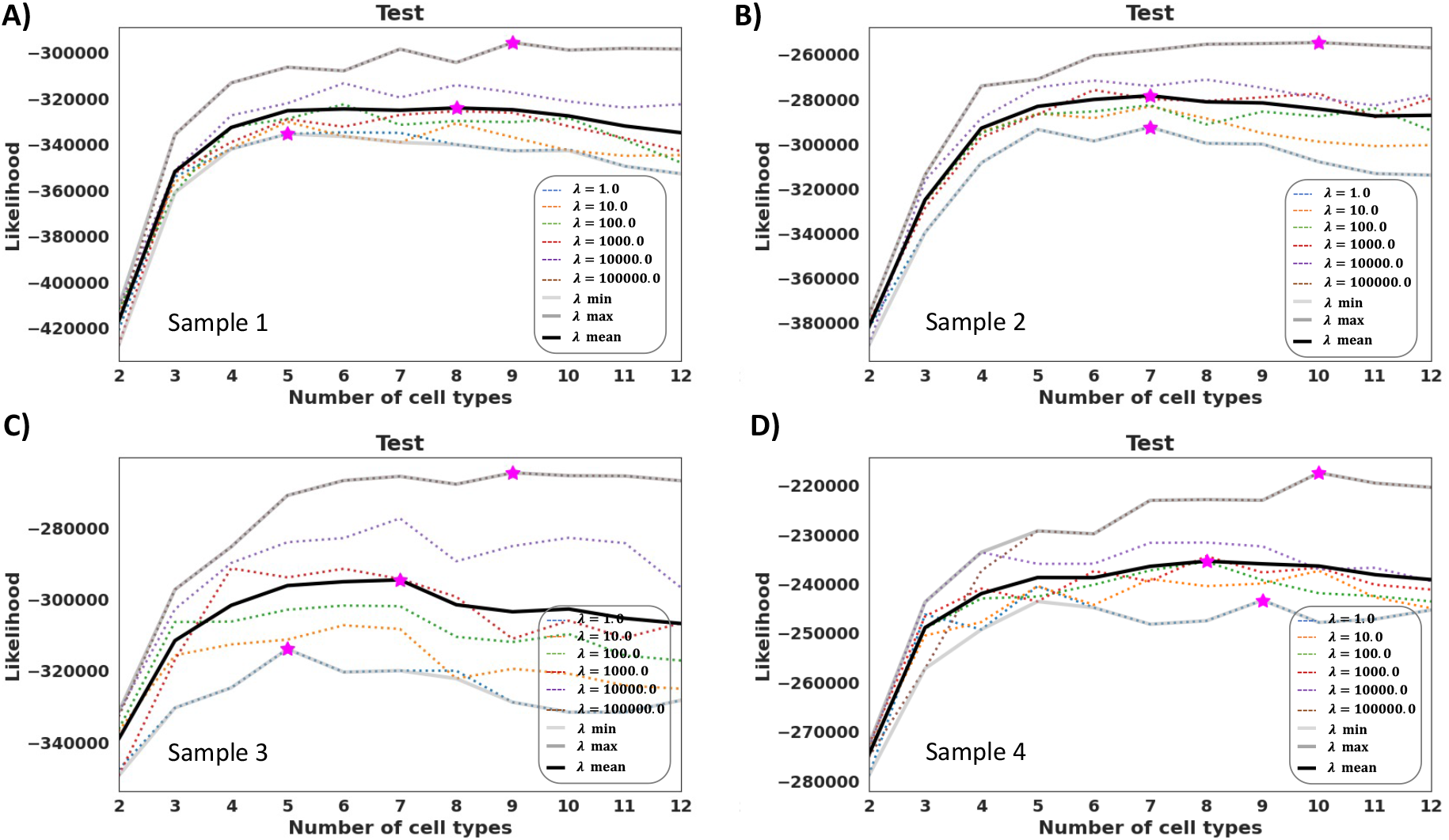
Inferring the number of cell types with **MUSTANG**. The cross-validation log-likelihood for different values of *λ* as well as the average log-likelihood across different *K* values are shown for four different bone tissue samples.

**Figure S3:**
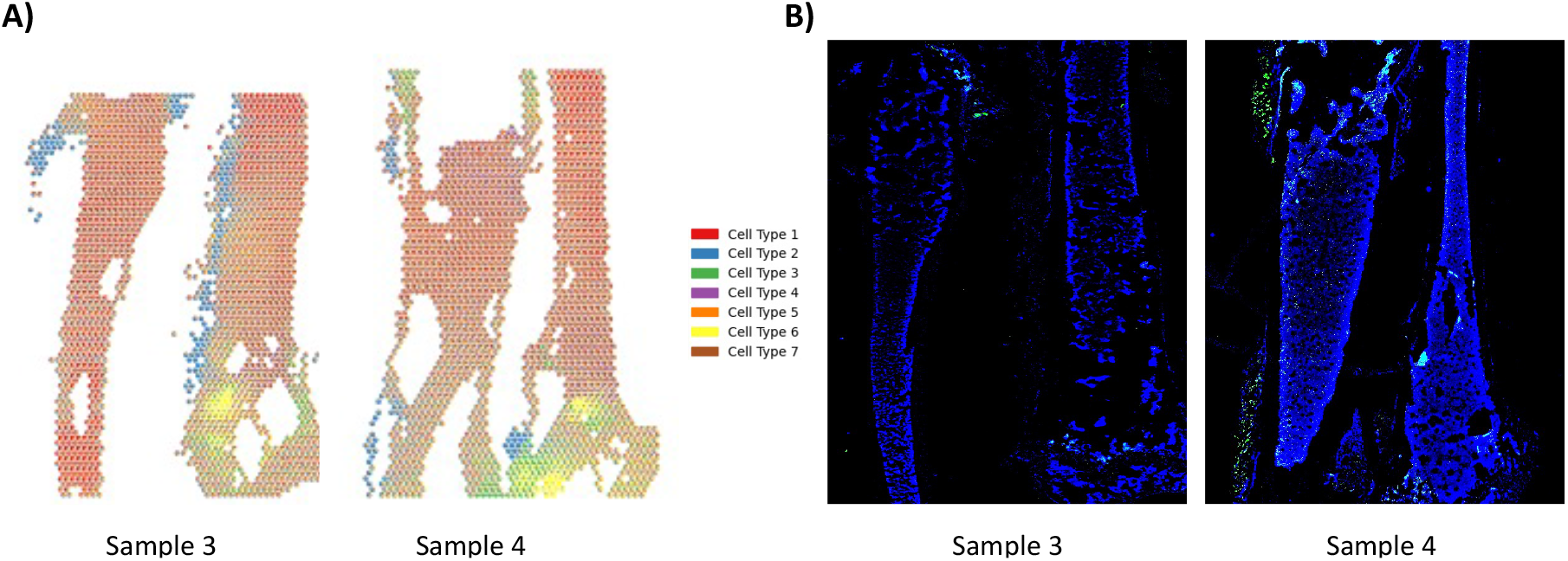
Additional results of jointly analyzing mouse bone marrow samples with **MUSTANG**. A) Spot-based spatial pie charts of **MUSTANG**-inferred cell type proportions for (left) sample 3 and (right) sample 4. B) Matching **IF** staining images of (left) sample 3 and (right) sample 4.

